# Cortical subnetworks encode context of visual stimulus

**DOI:** 10.1101/452219

**Authors:** Jordan P. Hamm, Yuriy Shymkiv, Shuting Han, Weijian Yang, Rafael Yuste

## Abstract

Cortical processing of sensory events is significantly influenced by context. For instance, a repetitive or redundant visual stimulus elicits attenuated cortical responses, but if the same stimulus is unexpected or “deviant”, responses are augmented. This contextual modulation of sensory processing is likely a fundamental function of neural circuits, yet an understanding of how it is computed is still missing. Using holographic two-photon calcium imaging in awake animals, here we identify three distinct, spatially intermixed ensembles of neurons in mouse primary visual cortex which differentially respond to the same stimulus under separate contexts, including a subnetwork which selectively responds to deviant events. These non-overlapping ensembles are distributed across layers 2-5, though deviance detection is more common in superficial layers. Contextual preferences likely arise locally since they are not present in bottom up inputs from the thalamus or top-down inputs from prefrontal cortex. The functional parcellation of cortical circuits into independent ensembles that encode stimulus context provides a circuit basis underlying cortically based perception of novel or redundant stimuli, a key deficit in many psychiatric disorders.

**One Sentence Summary:** Visual cortex represents deviant and redundant stimuli with separate subnetworks.

---

In a healthy neocortex, neural responses to incident sensory stimuli are modulated by past experience on short time scales (.01-10 sec) as well as long (>10 sec). For instance, repetition of a given stimulus at a rate of .1-2 Hz results in a phenomenon termed “stimulus specific adaptation” or SSA, wherein response magnitudes decrease rapidly to the repeated or “adapted” stimulus(*1*–*3*). In contrast, when a stimulus is unexpected given the established contextual regularities (e.g. probability of occurrence), a phenomenon termed “deviance detection” is observed, wherein response are augmented beyond the typical magnitude observed in a neutral context (*1*).

These phenomena have been well studied at either the single neuron level in animals (*1*–*3*) or at the gross cortical level with EEG or MEG in humans (*4*–*6*). Resulting work highlights a role of NMDA-receptors (*7*), cholinergic modulation (*6*), and local somatostatin interneurons (*1*, *3*). Such findings are clinically relevant since SSA and deviance detection, and the related “mismatch negativity” biomarker, are reduced in individuals with psychotic disease (*5*), providing the promise of linking specific neuropathologies with non-invasive biomarkers measurable in a clinical setting (*1*, *8*). This goal remains nevertheless difficult, perhaps due to an incomplete understanding of the fundamentals of cortical computation. For instance, recent work with multicellular recording techniques highlight that cortical neuronal populations tend to activate in groups and/or sequences which repeat during rest, sensory stimulation, and cognitive tasks, potentially outlining the preferred activity patterns or attractor “vocabulary” of the neocortex (*9*–*13*). Such dynamics are not evident in single neuron recordings or in gross electrophysiology (LFP/EEG), yet new results indicate that a breakdown in these local cortical activity patterns may be a fundamental aspect of schizophrenia (*14*, *15*). Given that context processing (*16*), and namely the mismatch negativity biomarker (*17*, *18*), is reduced in schizophrenia, it is crucial that these functions and their components (SSA, deviance detection) be understood at the level of the neuronal ensemble.

To investigate this, here we recorded the activity of populations of cortical neurons in awake mice (Figure 1A; n=23 experiments from 15 mice; 3140 cells) while they viewed a classic visual oddball paradigm (100% contrast, full field square-wave gratings oriented 0 vs 90 or 45 vs 135 degrees; redundant stimuli 88.5% probability, deviant stimuli [opposite orientation] 12% probability; Figure 1B). Responses in the oddball paradigm were compared to a reversed run (e.g. deviant stimuli in first run become redundant stim in second run 5 minutes later) and to a “many standards control” (*1*), wherein 8 different orientations were presented at the same rate with 12.5% probability, such that stimuli were neither redundant nor contextually deviant. We first employed fast two-photon calcium imaging (30Hz resonance scanning) with GCaMP6s in layer 2/3 of V1, focusing on increases in fluorescence (Figure 1C,D), to quantify neuronal responses in the same neurons to the same stimulus under three 3 separate contexts (redundant, deviant, neutral). Analyses focused on robust visually driven neurons showing Z-scored average responses greater than 3 stds above baseline to one of two orientation stimuli in at least one of 3 contexts (neutral, redundant, deviant; n=445 neurons, about 28% of total cells).

**Fig. 1.**
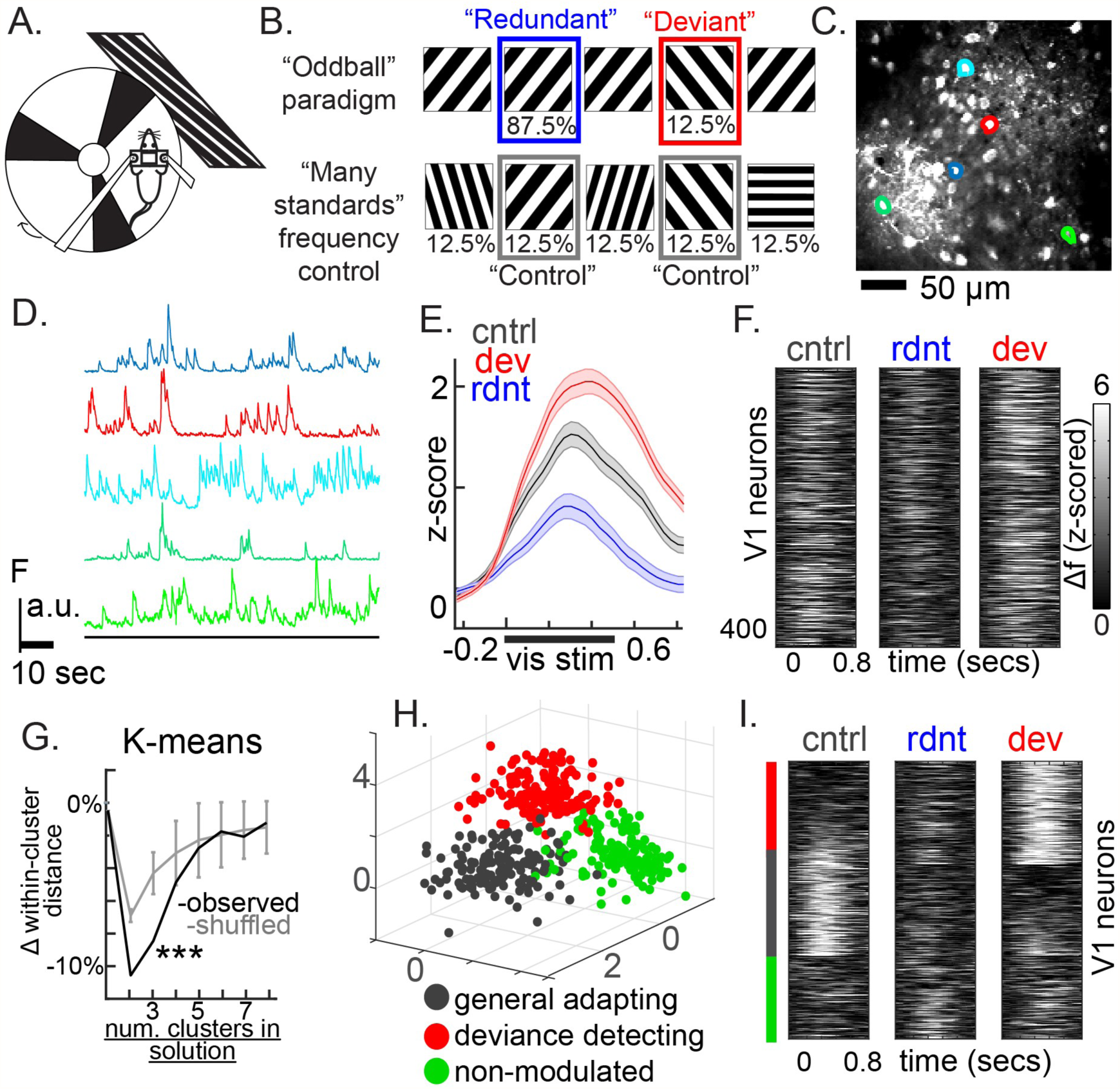
Contextual modulation of subnetworks of neurons. (A) Head-fixed awake mice viewed full-field visual grating stimuli in (B) a typical oddball paradigm and a many-standards control run. (C) Two-photon calcium imaging quantified (D) neural activity in layer 2/3 of V1 cortex. (E) Averages across all visually driven neurons indicate deviance detection (dev>cont) and stimulus-specific adaptation (cont>rdnt) at the population level. (F) Unsorted average responses for individual neurons to the same stimulus across three contexts. (G) k-means analysis indicates that the three cluster solution, but not higher, reaches significance (p<.001; error bars=3.10 standard deviations from mean). (H) The average responses of individual neurons plotted in 3-dimensional space (X=rdnt, Y=cntrl, Z=dev) show 3 spatially distinct clusters, which are evident in (I) the sorted responses from (F). These include a deviance detecting cluster (top; red), a “general adapting” cluster (middle; gray), and a non-modulated/mixed cluster (bottom; green).

When averaged across all visually driven cells in layer 2/3, cortical responses were suppressed to redundant stimuli (SSA; focusing on third redundant stimulus in the sequence; cont>rdnt; t^paired^(444)=3.94, p<.001; Fig 1E) and amplified to deviant stimuli (deviance detection; dev>cont; t^paired^(444)=3.67, p<.001; Fig. 1E). The gross local field potential (LFP), current source density (CSD) estimate, and time-frequency decomposition of the LFP confirmed the presence of SSA and deviance detection present across layers (Fig. S1). While the population averages showed SSA and deviance detection, it is possible that not all neurons were equally affected. Indeed, examination of individual neuronal responses across neutral, redundant, and deviant contexts suggested variability in the degree to which individual neurons expressed both SSA and deviance detection (Figure 1F). We hypothesized that this variability may show taxonicity, or, in other words, that distinct sub-populations of neurons could be carrying out deviance detection vs SSA to different degrees. If so, the population average, and LFP measurements, could exhibit both but due entirely to separate mixed populations in the local cortical area. To explore this, K-means clustering analyses was employed on averaged neuronal responses to each visual stimulus context, holding individual neuron/stimulus combinations as individual observations (445 neurons, 3 dimensions: redundant vs deviant vs control). This was carried out systematically, constraining the k-means solution from 2 through 8 clusters on the main dataset (see Methods). The analysis suggested that 3 functional clusters of context processing subtypes were present (Figure 1H). These subnetworks included a “general adapting” subset (33% of cells), whose responses were largest in the control paradigm, yet suppressed during the oddball paradigm regardless of whether the preferred stimulus was redundant or deviant. The second ensemble was a “deviance detecting” ensemble (39% of cells), whose responses were mostly silent in neutral or redundant contexts, but were largely amplified when its preferred stimulus was contextually deviant. The third ensemble (28% of cells) appeared to show no apparent contextual modulation, expressing equal response magnitudes to a visual stimulus when it was redundant, deviant, and neither (Figure 1I; S2A-C). Importantly, these clusters replicated with two separate validation approaches, and cells within the same clusters showed stronger correlations across the imaging session than cells in separate clusters (See supplementary Text and Fig. S2).

Next we sought to determine the degree to which context preferring subnetworks were present in different cortical layers. We employed a holographic imaging approach [Fig 2a, (*19*, *20*)] which enabled fast volumetric calcium imaging of large populations of neurons (400-2000 cells) across 500um of depth (5 mice, 10 experiments). GCaMP6s (a green calcium indicator) was expressed in neurons in layers 1 through 3 (depth 1 to 350um), while jRGECO1b (a red calcium indicator) was expressed in neurons in layers 4 and 5 (depth 350-550um; Figure 2b,c). Imaging was carried out with two lasers (920nm and 1064nm), which simultaneously scanned two different depths (layer 1~3, and 4~5 respectively) using a resonant scanner. The signal was separated by the emission color of GCaMP6s and jRGECO1b and an electrically tunable lens and a spatial light modulator were implemented in the two beam paths to enable fast sequential scanning of different focal planes (30~40um plane separation across 150 to 530um depth range; Fig 2a). A total of 10 imaging planes were imaged at a 10Hz volume rate (figure 2b). Images were separated into layers (2-5) based on laminar boundaries and pre and post-imaging z-sectioning, and later confirmed with histology.

**Fig. 2.**
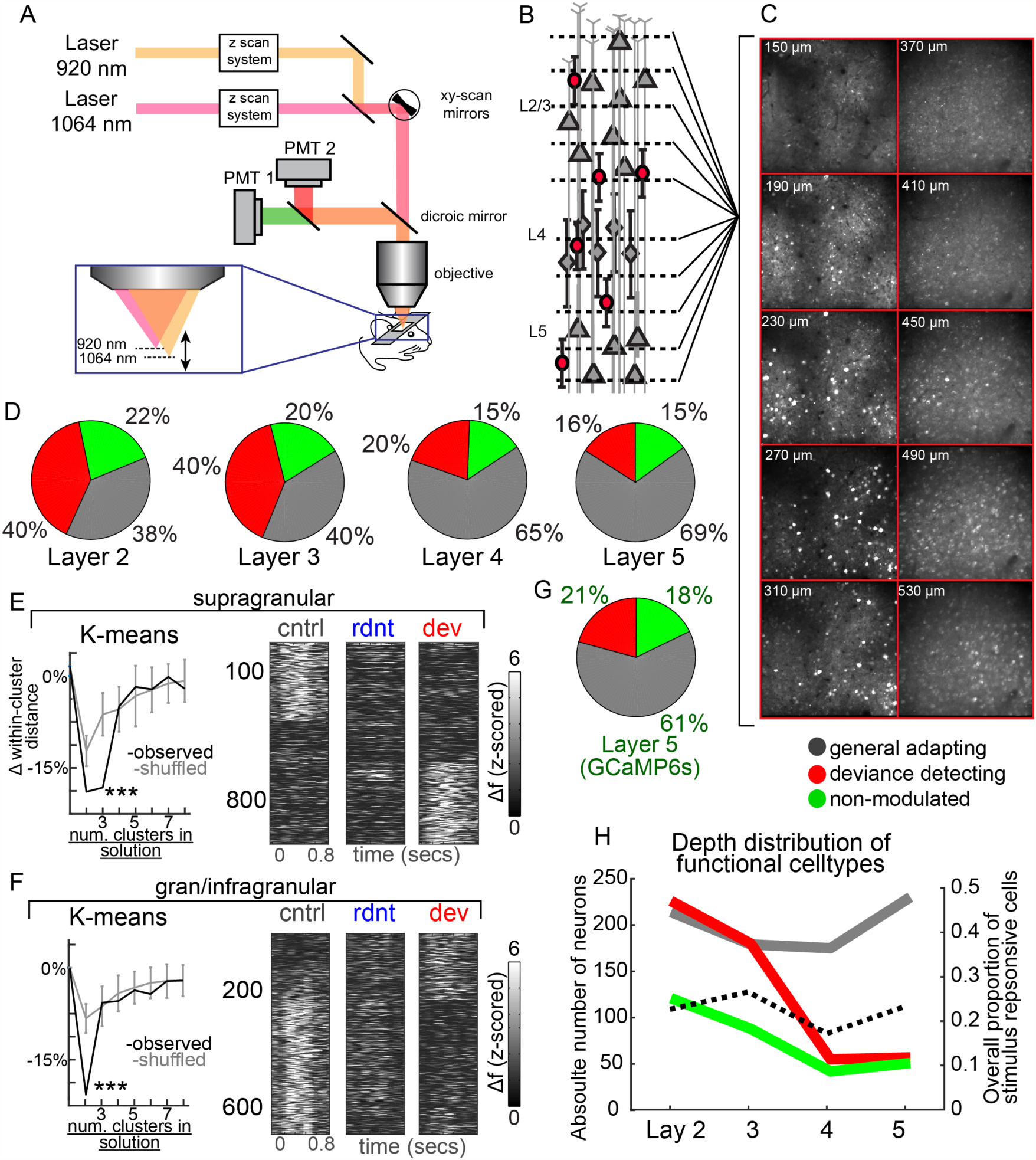
Context processing subnetworks differ across cortical layers. (A) Schematic of microscope setup involving two-excitation lasers with distinct z-sectioning strategies (SLM and ETL). (B) This enabled simultaneous imaging of neurons across 10 planes and ~400um of cortical depth. (C) Average images from one experiment at each depth. (D) Relative distributions of three context preference ensembles from 4 subsections of cortex. (E) K-means analysis suggests the presence for 3 clusters for supragranular populations (140-310um) which show similar functional properties and distributions as initial experiments, while (F) gran/infragranular subpopulations (370-530um) exhibit only 2 clusters, with a majority showing general adapting properties. (G) These superficial vs deep differences are not due to the calcium indicator used, as GCaMP6s expressing cells show the same distribution as jRGECO1b. (H) Overall proportion of responsive neurons was not different across layers (dotted line), although sub-cluster proportions varied significantly.

Neurons in all layers showed robust visually driven neurons (see above). We first focused analyses layers 2-5: 150-230um (layer 2; 561 out of 2474 cells responsive, 23%), 270-310um (layer 3; 447/1683, 26%), 370-410um (layer 4; 310/1580, 20%), and 450-530um (layer 5; 338/1450, 23%). Layers did not significantly differ in the proportion of visually responsive cells within mice (*F(3,3)*=3.12, *p*=.08). When the 3 cluster K-means solution was applied to each of the 4 laminar sections, the results revealed a surprising difference in relative distribution of cell response types (Fig 2D). Layers 2 and 3 displayed the same relative distribution of general adapting, deviance detecting, and non-modulated cell responses as the previous set of mice in the previous setup, replicating the initial results (Fig. 1). On the other hand, layers 4 and 5 displayed about 50% more “general adapting” cells, and 50% fewer “deviance detectors”. Based on similarities in cluster distributions, we split these two subsets of cells into two populations: supragranular and gran/infragranular. This confirmed the statistical difference in cluster distributions (Fig 2D; *X*^*2(supra-vs-gran/infra)*^*(2)*=16.74, *p*<.001). We re-computed the number of clusters using the shuffling approach described above. Indeed, supragranular population analysis suggested 3 clusters, as before, which showed the same activity profiles as the original three clusters (Fig. 2E). On the other hand, the gran/infragranular population analysis suggested only two clusters (Fig. 2F, left), with the vast majority (about 80%) displaying general adapting properties, and only about 20% showing deviance detection. This difference was not due to the difference in calcium indicator (GCaMP6s vs jRGECO1b), since expressing GCaMP6s at 500um depth in two mice replicated these effects (Fig. 2G; n=52 responsive cells, 2 mice; *X*^*2(gcamp-vs-jrgeco1a)*^*(2)*=0.78, *p*=.67). Assessing this result in terms of absolute number of neurons suggests that deviance detectors in particular are almost four times more common in superficial layers than in deep layers, while other subtypes remain mostly the same (Fig 2H).

We next explored whether aspects of context processing were present in bottom-up inputs to the visual cortex from the dorsal lateral geniculate nucleus of the thalamus (dLGN). The dLGN is the primary relay nucleus for visual information from the retina to the cortex(*21*). We expressed GCaMP6s in the dLGN (Fig. 3A; S3A) and performed two-photon calcium imaging of the axonal boutons from dLGN cells in layers 1-4 in the V1 (Fig 3B; (*22*)). Interestingly, at the population level, inputs to V1 showed clear reductions in activity to redundant stimuli (SSA; *t*(97)=6.01, *p*<.001), but did not show amplified responses to contextually deviant stimuli (no significant deviance detection; *t*(97)=.617, *p*=.53; Fig. 3C). Further, a cluster analysis indicated that, unlike V1 layer 2/3 neurons, only 2 functional clusters were present among thalamic inputs to V1 (Fig. 3D), and neither displayed deviance detection (Fig. 3E). The biggest cluster (70% of inputs), displayed basic SSA, but equivalent responses to neutral vs deviant stimuli. The remaining 30% showed no contextual modulation (Fig. 3F,G,H). This provides strong evidence that deviance detection present in primary visual cortex is not inherited in a bottom-up, feedforward manner from the thalamus. Further, it suggests that some aspects of SSA are inherited from the thalamus, as demonstrated by others (*3*).

**Fig. 3.**
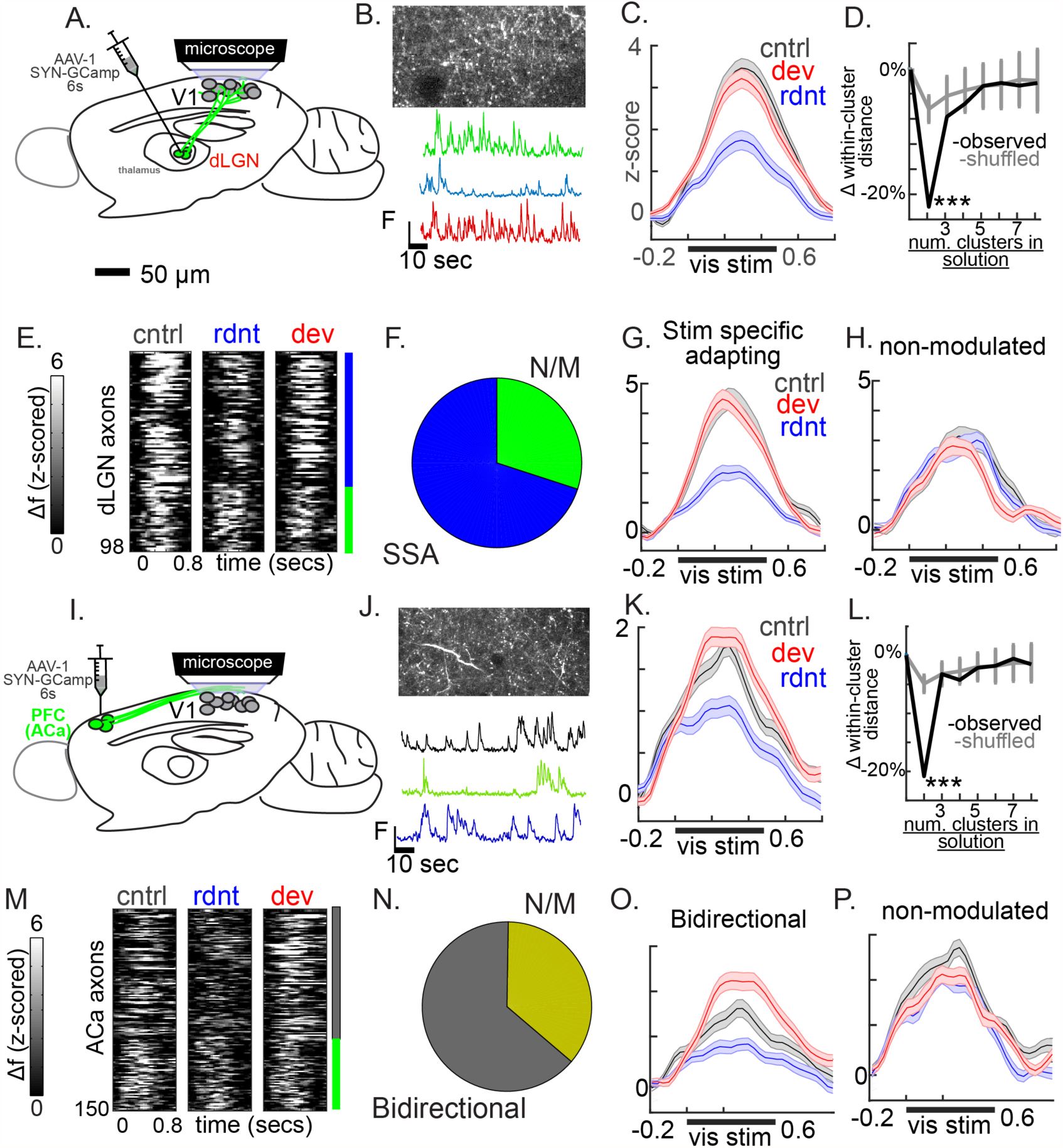
Context processing in bottom-up and top-down inputs to V1. (A) Schematic of viral injection to dLGN of thalamus, and (B) GCaMP6s expression in dLGN axons in V1 (350um below dura) shown in an average projection (above) and 3 example traces from boutons. C) Average responses from responsive cells. (D) Results of K-means analysis vs shuffled surrogate, suggesting 2 clusters. (E) Individual axonal averages, sorted by their clusters represented in (F) a pie chart showing their relative proportions and (G, H) line plots showing their group averages to visual stimuli. (I-J) display the same plots for PFC (Cg1) axonal inputs to V1.

Mouse V1 also receives top-down inputs from executive brain regions, like prefrontal cortex area Cg1 [anterior cingulate area; (*23*)]. These inputs have a net inhibitory effect which aid in simple visual discrimination tasks (*24*), yet their role in providing context information to sensory areas, particularly in the processing of deviant vs redundant events in the oddball paradigm, is unknown. This information could be critical for relating known deficits in context processing (e.g. mismatch negativity) to prefrontal cortical and cognitive dysfunction known to be present in individuals with schizophrenia. Using a similar axon imaging approach as described above, we imaged the activity of PFC inputs to layer 1 of V1 (Fig. 3I,J; S3B). Like thalamic inputs to V1, PFC inputs to V1 displayed SSA (*t*(149)=.4.17, *p*<.001) but not deviance detection (*t*(149)=1.162, *p*=.24) when considered *en masse* (Fig. 3K). Further, only 2 functional clusters were present among PFC to V1 (Fig. 3L,M). The biggest cluster (64% of inputs), was bidirectionally modulated, showing both SSA and deviance detection. The remaining 36% showed no contextual modulation (Fig. 3N, O, P). So while these top-down inputs appear to show different dynamics at the population level and a different make-up of sub-populations than V1, it still remains possible that PFC inputs contribute to deviance detection and context processing clusters in V1. For instance, bidirectionally modulating PFC inputs could primarily target deviance detecting neurons in V1, amplifying their properties.

In order to determine whether PFC inputs to V1 play a causal role in V1 context processing in the oddball paradigm, we expressed an optogenetic silencer, ArchT, in prefrontal cortical neurons under the synapsin promoter (Fig 4A). We focused a 4mW 617nm LED directed with a cannula focused with a ~1 radius centered over a 2mm craniotomy over the visual cortex (Fig 4A) and suppressed axons (Fig. 4B) from PFC. We confirmed this effect by tracking neural activity in PFC axons with GCaMP6s while suppressing them with coexpressed ArchT and 2 second LED pulses while performing 2P calcium imaging (Fig. 4C,D). Indeed, after photostimulation, axons showed a dramatic decrease in calcium transients (averaged over cells and 20 trials, Fig. 4D). Further, we recorded local field potentials with 16-channel multielectrode arrays while suppressing axons from PFC locally with ArchT (Fig. S4a; n=7) and compared this with recordings from mice expressing only GCaMP6s in PFC axons (Fig S4b; n=6; LED only controls). This technique showed that, despite focusing the LED over V1, there was a visually evoked potential present in the current source density plots even when ArchT was not expressed (Fig. S4B). Nevertheless, the ArchT induced current i) started immediately (<6ms) after the onset of the LED, while the visually induced response started 60ms later, ii) had a distinctly different current source distribution (difference plot Fig. S4C) and iii) was much stronger in superficial layers (Fig S4D), perhaps due to a combination of the fact that PFC axons terminate mostly in layer 1 and 6 (*24*) and the fact that the LED illumination was likely strongest in superficial tissue due to scattering. While interpreting this LFP/CSD distribution is complex, it is clear that the effect of ArchT stimulation in V1 was present, dramatic, and sufficient to suppress PFC axon activity.

**Fig. 4.**
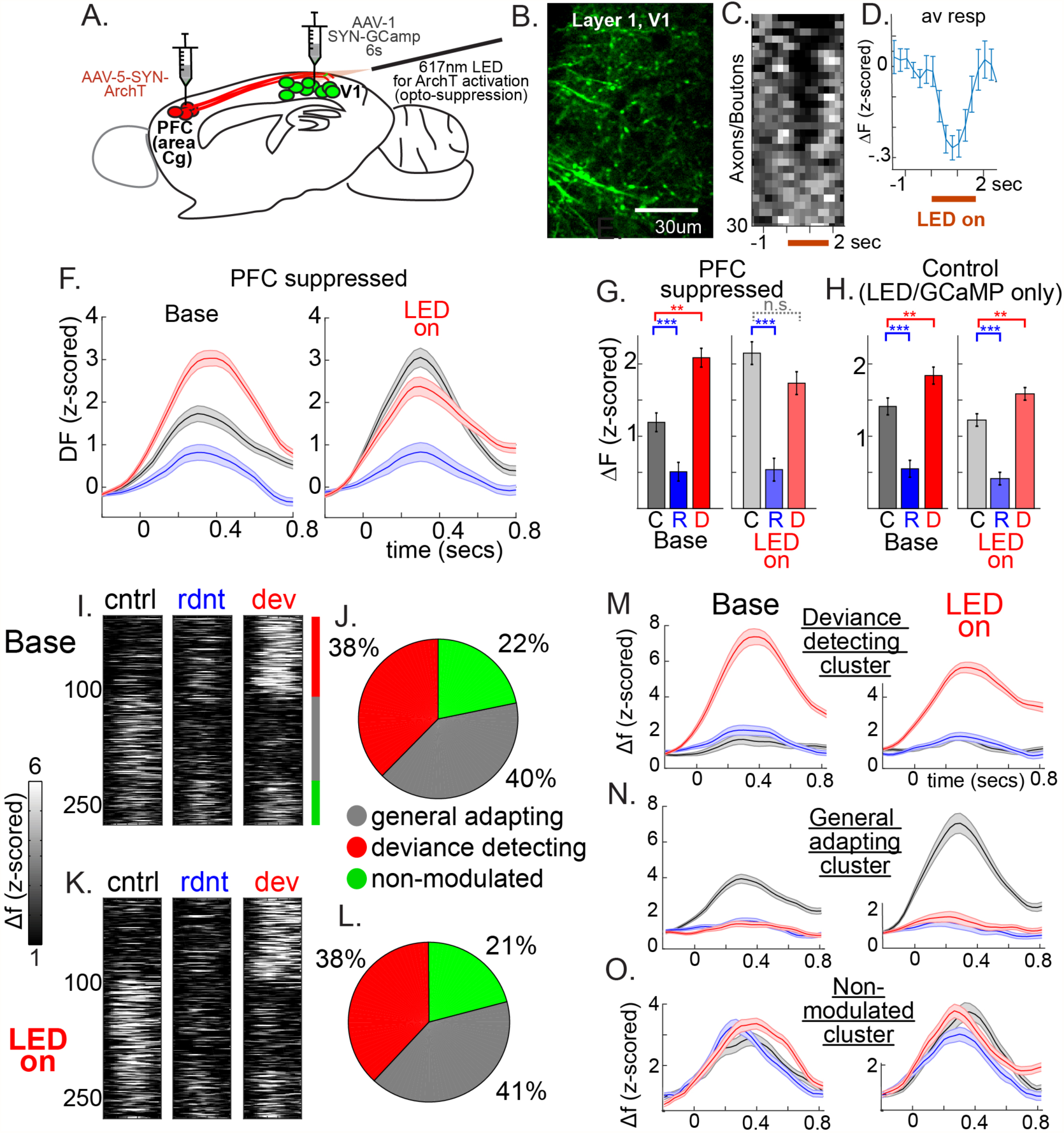
PFC influences on context processing in V1. (A) Schematic of viral injections to Cg1 of PFC and V1, and (B) GCaMP6s expression in PFC axons in V1 (50um below dura). (C) Average responses from axon segments expressing ArchT during LED stimulation. (D) axons from (C) averaged together. (F) Average responses across 317 V1 neurons before (left) and during PFC axon suppression. (G) Activity from stimulus interval from (F). (H) Same as (G) but from neurons from control mice (i.e. no ArchT). (I) pre-PFC suppression k-means sorted neuron average responses to stimulus types, and (J) relative proportion of each subcluster. (K,L) same as (I) and (J) but during PFC suppression. (M,N,O) averages across all neurons in each cluster before (left) and during PFC suppression. (**p<.01, ***p<.001).

We then recorded soma from V1 neurons expressing GCaMP6S in the oddball and control paradigms as described above. Then we repeated the photostimulation sequence, delivering 3.5mW of LED power over V1 every odd trial in mice expressing ArchT in PFC neurons (9 experiments in 7 mice). PFC suppression had a significant effect on SSA (F^interaction^(1,582)=4.21, p<.05) and deviance detection (F^interaction^(1,582)=7.19, p<.01). At baseline, neurons displayed clear SSA (*t*(313)=4.76, *p*<.001) and deviance detection (*t*(313)=3.95, *p*<.001), yet during runs with PFC suppression, SSA was still clearly present (*t*(269)=5.77, *p*<.001) but deviance detection was absent (*t*(269)=1.29, *p*=.20). This was driven mainly by an increase in the average responses to control stimuli (*t*(582)=2.68, *p*<.01), while responses to deviant (*t*(582)=-1.36, *p*=.17) and redundant stimuli remained the same (*t*(582)=1.04, *p*=.30). We repeated this in mice with only GCaMP6S injected as a control set (i.e. LED/GCaMP only; 15 experiments in 8 mice; Fig. 4H, S5), and such experiments showed no effect on SSA (F^interaction^(1,490)=0.00, p=.99) or deviance detection (F^interaction^(1,490)=1.47, p=.23). This same effect of increased response to neutral context stimuli, but not deviant stimuli was replicated with an entirely separate estimate of local neuronal firing, multiunit spiking from multielectrode recordings, in a separate set of mice (n=6; Fig S6; 100-200ms post-stim, dev vs cntrl; F(1,5)^interaction^=6.18, p<.05; F(1,5)^pre_dev_vs_cntrl^=10.61, p<.05; F(1,5)^post_dev_vs_cntrl^=.42, p=.54).

Finally, we explored whether PFC suppression affected the organization and context preference of local neuronal ensembles. Interestingly, three clusters were present before and during PFC suppression, and the proportion and nature of these cell sets were identical to the initial experiments and to each other, including general adapting (40-41%), deviance detecting (38%), and non-modulated (21-22%) subpopulations (Fig. 4I-O). Thus, PFC suppression appeared to eliminate deviance detection in the gross, unsorted output of V1 as evidenced by the overall calcium imaging grand average (Fig 4F,G) and by the multiunit activity (Fig S6). Yet “general adapting”, “deviance detecting”, and “no-modulation” subnetworks of V1 cells remained present and functionally distinct. Thus these context preferences may be a fundamental property of local layer 2/3 cortical circuits of visual cortex, perhaps “soft-wired” via modulatory interneuron effects (*25*) or even “hard-wired” (*26*). This idea is supported by these PFC suppression findings, along with the fact that dLGN inputs do not display deviance detection or similar context-preference clustering, and along with previous demonstrations of non-overlapping feedforward (layer 4-2/3) and interneuronal networks in local neocortex (*27*, *28*). Exhaustively ruling out the role of non-V1 regions in generating these distinct context-preferring clusters remains a future goal.

The discrepancy between effects present on the population average vs. ensembles is accounted for by the fact that the effects of PFC suppression on the context preferring ensembles individually was varied, with general adapting cells showing enhanced responses in their preferred context (i.e. control; *t*(236)=3.58, *p*<.001), while deviance detectors (responses to deviants; *t*(218)=-1.58, *p*=.14) and non-modulating cells showing no significant changes (average of all contexts; *t*(124)=0.53, *p*=.59). These non-ovelapping subnetworks could be hardwired or, alternatively, arise from synaptic plasticity as specific circuits crafted from a common neural network, as needed. The exact circuit mechanisms which account for this effect are unknown, perhaps involving inhibitory interneurons (*1*, *3*, *29*), but also potentially involving specific subcellular (e.g. proximal vs distal), cell-specific (e.g. interneuron subtypes), and/or layer specific targeting (e.g. layer 4 vs 2/3) of these two specific sub-populations arriving from PFC (bi-directional modulators and non-modulators). Future work will require targeting opto-or chemico-genetic tools to cells based purely on their functional properties during this paradigm, and/or cellular reconstructions from cells recorded *in vivo*.

In summary, our results reveal a functional parcellation of cortical circuits into specific ensembles of cortical neurons engaged in different computations with the same inputs. These distinct context preferring subnetworks are present in primary sensory cortex and provide a circuit phenomenology underlying how the brain responds during the “oddball” paradigm, a paradigm of great importance for basic psychology and understanding the pathophysiology of major psychiatric disorders (*4*).

## Acknowledgments

The authors would like to acknowledge Dr. Reka Letso for technical contributions and comments and numerous members of the Yuste lab for comments.

## Author contributions

Conceptualization, J.P.H., Y.S., R.Y.; Methodology, J.P.H., W.Y., Y.S.; Investigation, J.P.H.; Y.S., S.H.; Data curation: Y.S., S.H.; Formal analysis, J.P.H; Writing–Original Draft, J.P.H.; Writing–Review & Editing, J.P.H, Y.S., S.H., W.Y., R.Y.; Project administration: R.Y. Funding Acquisition: R.Y., J.P.H.

## Data and materials availability

All data is available in the main text or the supplementary materials.

## Supplementary Materials

Materials and Methods

Figures S1-S6

References (*30-43*)

